## Supplemental figures for "Local synchronization of cilia and tissue-scale cilia alignment are sufficient for global metachronal waves"

FIGURE S1: **'Quantification of multiciliated cell features.'**

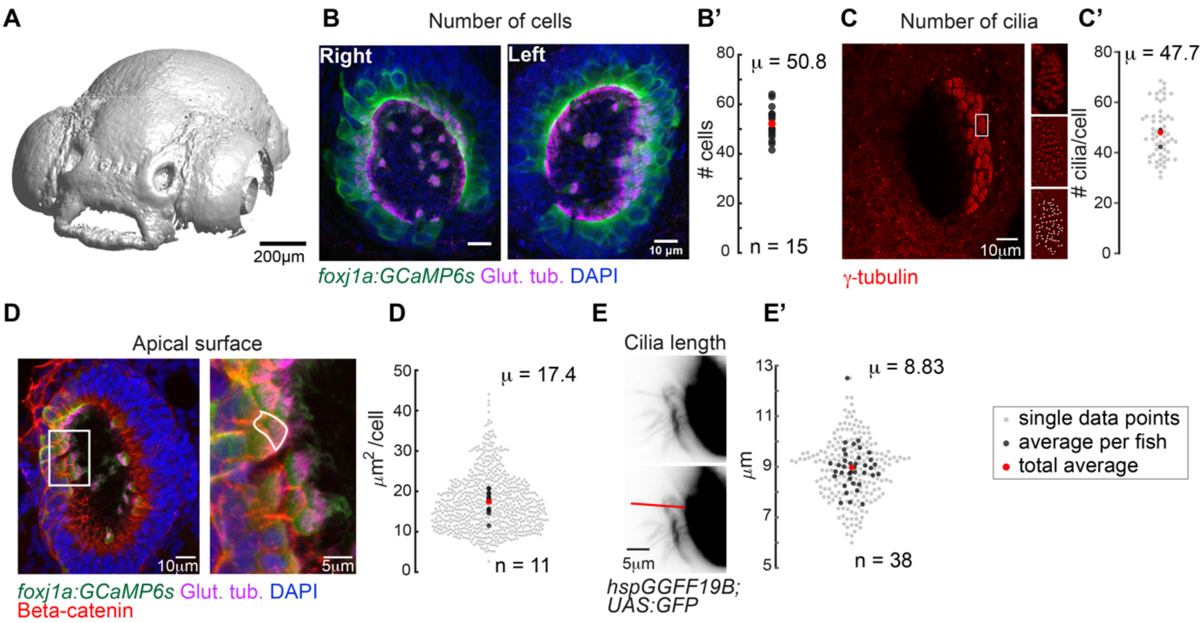


*(A) Surface rendering of a zebrafish head at 4dpf (using a transgenic lines expressing Cherry in all cells, ubi:zebrabow). Scale bar is 200µm. (B) A representative example of a left and right 4dpf zebrafish nose. Motile cilia are in magenta (glutamylated tubulin), nuclei in blue (DAPI), and multiciliated cells in green (foxj1a:GCaMP6s). Scale bar is 10µm. (B') There is a total number of 50.8 cells per fish (± 6.24 SD; n = 15). (C) To quantify the number of cilia per cell, basal feet were immunostained with gamma-tubulin (red). Insets display for one representative cell the raw fluorescence (top), the signal after applying a bandpass filter (middle), and the detected peaks corresponding to each individual basal body (bottom) (C') a cell bears 47.7 cilia (± 9.94 SD; n = 4). (D) To quantify the apical surface of a multiciliated cells, (foxj1a:GCaMP6s) larvae expressing GFP in multiciliated cells were immunostained for cilia (glutamylated tubulin, magenta), nuclei (DAPI, blue) and cell borders (beta-catenin, red). Right: example of a manual tracing of apical surfaces (D') the apical surface spans 17.4µm$^2$ (± 6.31 SD; n = 11). (E) To quantify ciliary length, motile cilia expressing GFP (hspGGFF19B:UAS:GFP) were manually traced from light-sheet microscopy still images. Scale bars are 5µm. (E') cilia are 8.83 µm long (± 0.862 SD; n = 38). All n refer to the number of fish. In the scatterplots, the individual data points are light-grey, an average number per fish is in black, and a total average is in red.*

**FIGURE S2 'Distribution of beat frequencies.'**

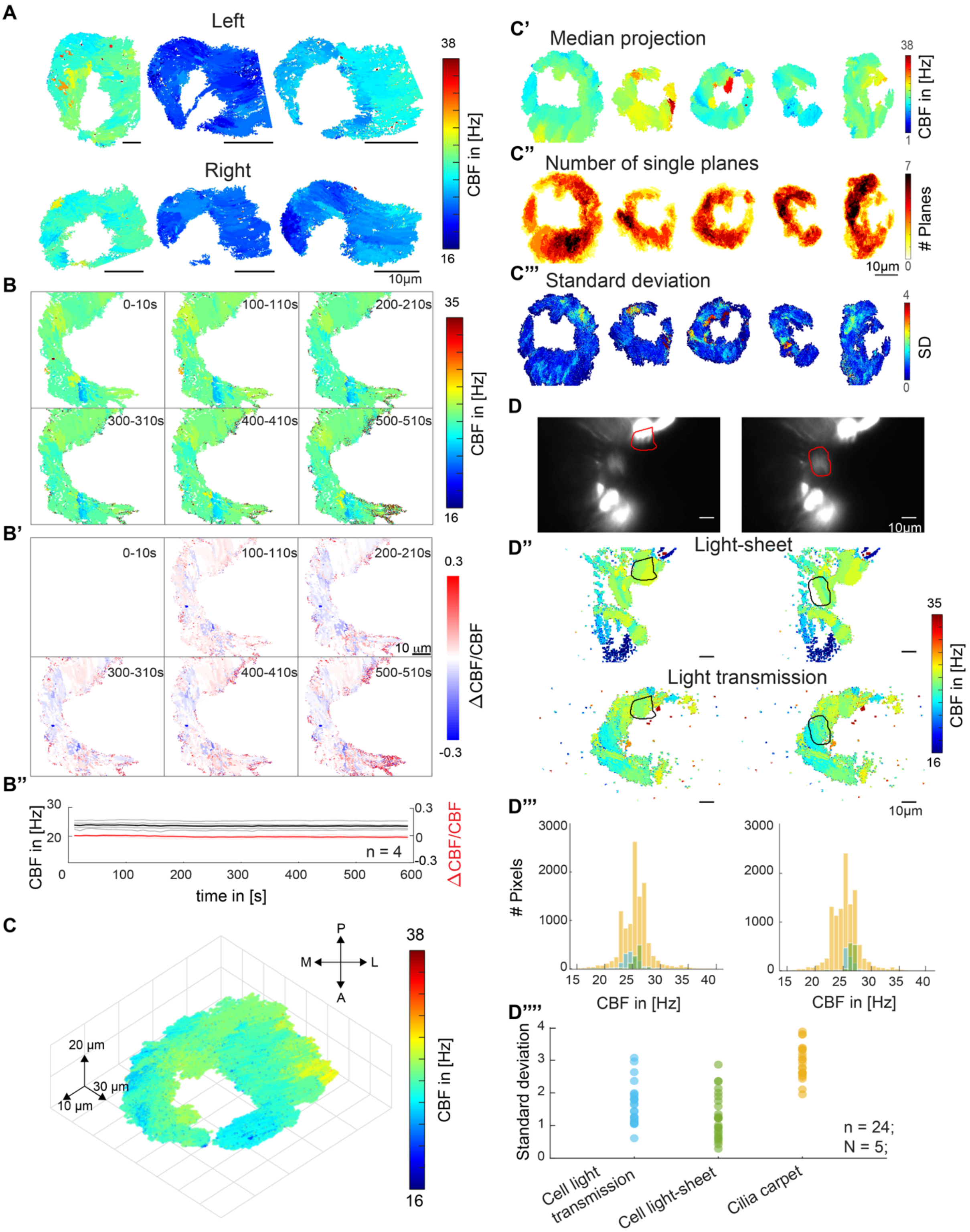


*(A) Ciliary beat frequency of 6 different fish, for left (top) and right (bottom) noses show high levels of heterogeneity. (B) Ciliary beating remains relatively constant over time. Ciliary beat frequency for a representative example over the course of ten minutes. Six instances are shown for frequency (top) and ΔCBF/CBF (B'). Values of four fish are depicted for both frequency and ΔCBF/CBF (B''). (C) A 3D CBF map across seven 10µm depth planes shows that frequency patches are consistent across Z planes. (C'-C''') Median and standard deviations for all 3D recordings. All scale bars are 10µm. (D) Cilia from individual cells beat*

*at a similar frequency. (D-D'') Representative examples of the beating of one cell versus the entire multiciliated epithelium. A hspGGFF19B:UAS:GFP animal recorded with light sheet microscopy (D), CBF in the light sheet (top), and CBF in light transmission (bottom). (D''') Corresponding CBF histograms show the entire frequency map (orange), together with the light sheet CBF (green), and light transmission CBF (blue) of individual cells. (D'''') Quantification of the standard deviation of all histograms show that standard deviation is smaller for individual cells than for the entire epithelium.*

**FIGURE S3: 'Ciliary beating in different viscosity conditions'**

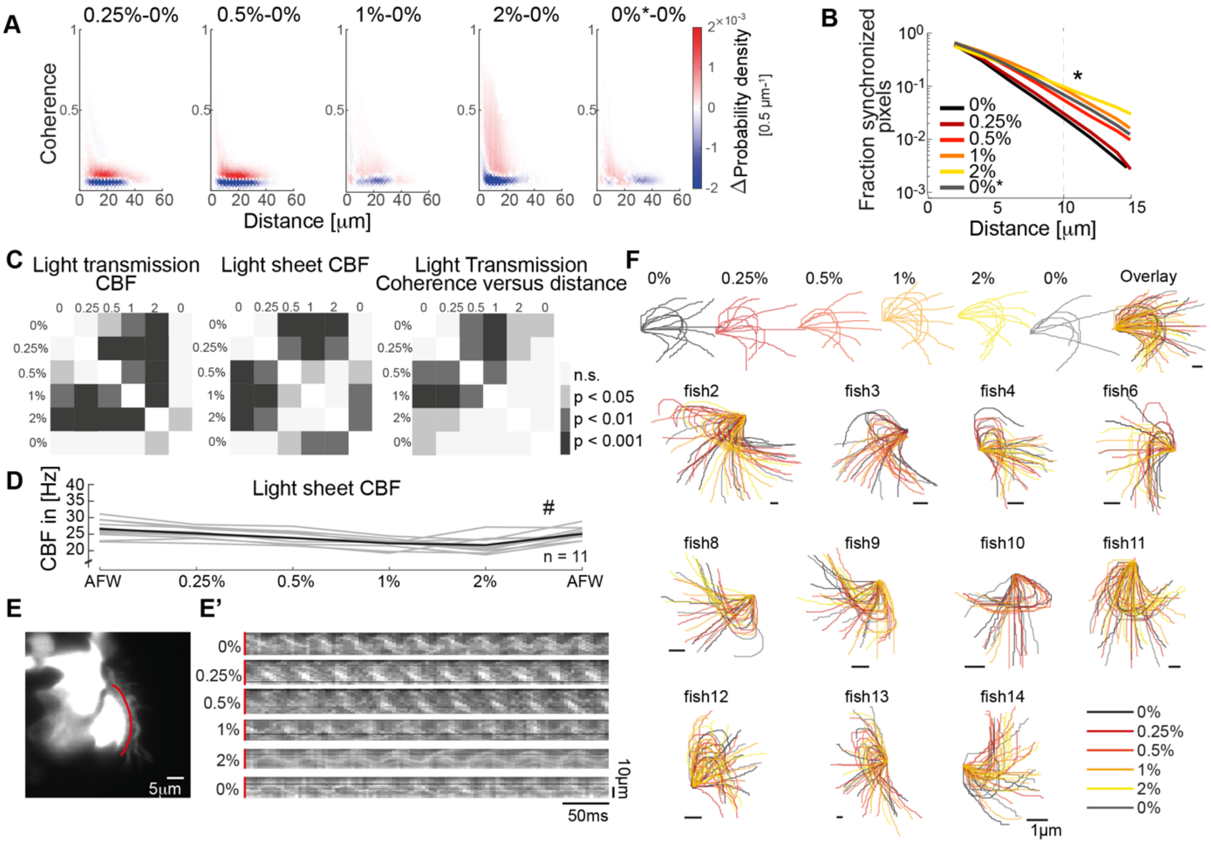


*(A) Difference probability histograms between the various viscosity conditions for the example shown in Fig 3C. (B) The fraction of synchronized pixels (coherence>0.25) increase with viscosity. Mean and standard deviation are shown. p = 3· $10^{-5}$ with ANOVA-N. (C) Significance tables depicting Tukey-Kramer post hoc multiple comparisons procedures for CBF in light transmission, CBF in the light sheet, and mean coherence versus distance measurements shown in Figure 3F. (D) Ciliary beat frequency of single cells decreases upon increasing viscosities as imaged by light-sheet microscopy. A repeated measures ANOVA (#) indicates a significant effect of viscosity conditions on CBF (p = 0.0011; n=11). (E) Representative light sheet recording of a hspGGFF19B:UAS:GFP zebrafish nose exposed to increasing concentration of methylcellulose shows changes in ciliary beating dynamics. (E') Kymographs (red) on transverse ciliary beating in recordings upon increasing viscosities. (F) Manually traced ciliary waveforms upon increasing viscosities (n = 11).*

FIGURE S4: **'Metachronal wave directions and lengths'**

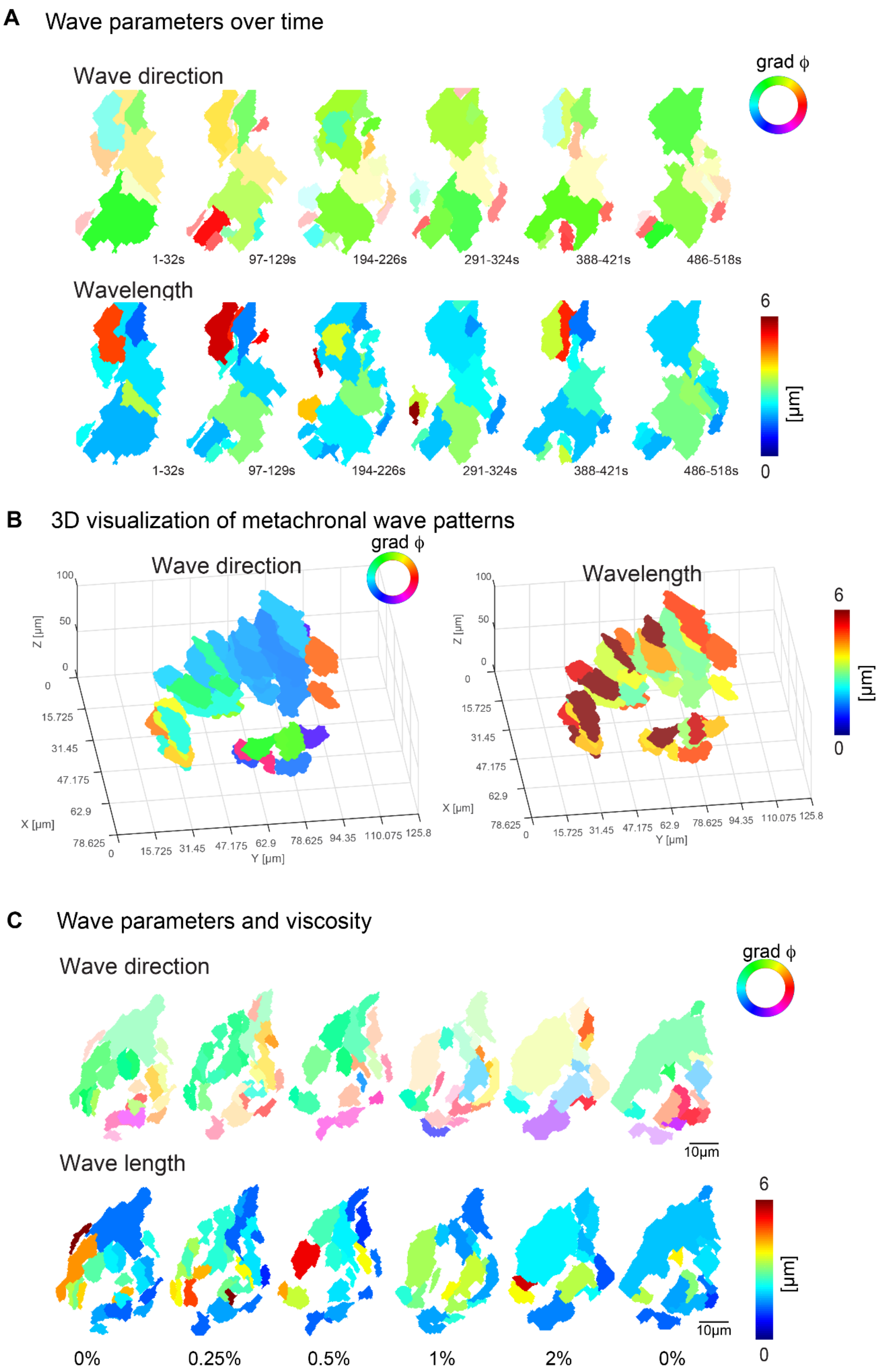

*(A) Wave parameters, wave direction (top), and wavelength (bottom), for a representative fish over the course of ten minutes. (B) wave direction (top), and wavelength (bottom), for a representative fish across seven 10μm depth planes (C) A representative example of wave directions (top) and wavelengths (bottom) upon increasing viscosities. Note that the transparency in wave direction reflects the inverse standard deviation.*

FIGURE S5: **'Metachronal wave and cilia beating directions'**

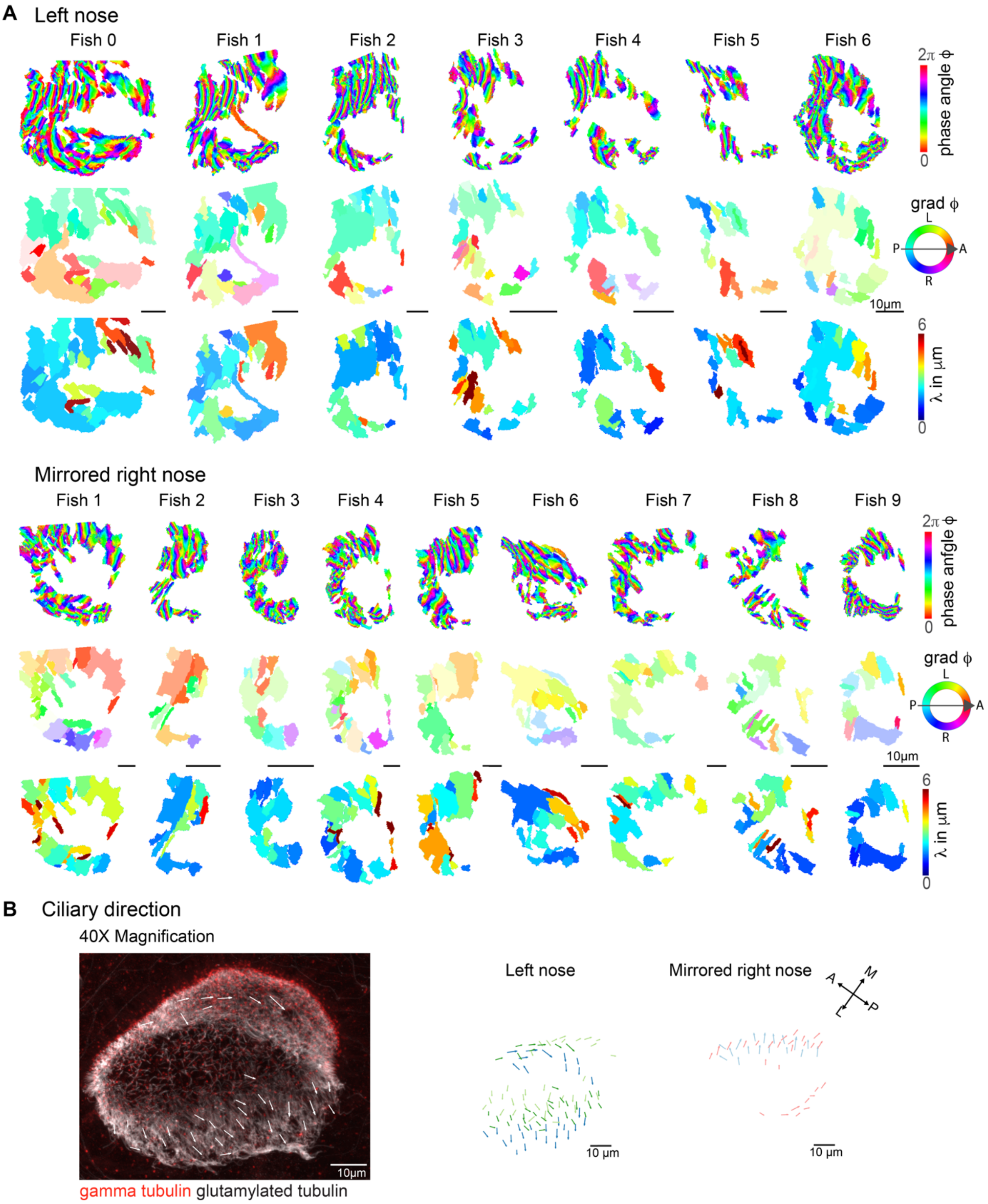

*(A) Wave parameters, phase angles, wave direction, and wavelength, for all aligned left (top) and right (bottom) noses. Note that the transparency in wave direction reflects the inverse standard deviation. (B) Immunohistochemistry on a 4dpf left zebrafish nose stained for gamma-tubulin (red) and glutamylated tubulin (white) at 40X. Zoom in on one cell displays how ciliary beat direction is determined (top). Overlay of the beat directions of multiple fish for left (n=3) and right noses (n=2) show identical results as with a 20X objective.*

**Video S1:** Measurement of metachronal wave properties in segmented frequency patch

**Video S2:** Measurement of metachronal wave in a 30s long recording. Each frame of the movie represents the analysis of 20s-long sliding windows. The timer indicates the center of the sliding window.

**Video S3:** Metachronal wave direction is stable over time (total of 10min) as shown for 4 examples. Every frame of the video corresponds to the output of a Fourier Transform calculated over a 30s timebin.
